# Major role of iron uptake systems in the intrinsic extra-intestinal virulence of the genus *Escherichia* revealed by a genome-wide association study

**DOI:** 10.1101/712034

**Authors:** Marco Galardini, Olivier Clermont, Alexandra Baron, Bede Busby, Sara Dion, Sören Schubert, Pedro Beltrao, Erick Denamur

## Abstract

The genus *Escherichia* is composed of several species and cryptic clades, including *E. coli*, which behave as a vertebrate gut commensal, but also as an opportunistic pathogen involved in both diarrheic and extra-intestinal diseases. To characterize the genetic determinants of extra-intestinal virulence within the genus, we carried out an unbiased genome-wide association study (GWAS) on 370 commensal, pathogenic and environmental strains representative of the *Escherichia* genus phylogenetic diversity and including *E. albertii* (n=7), *E. fergusonii* (n=5), *Escherichia* clades (n=32) and *E. coli* (n=326), tested in a mouse model of sepsis. We found that the high-pathogenicity island (HPI), a ∼35 kbp gene island encoding the yersiniabactin siderophore, is highly associated with death in mice, surpassing other associated genetic factors also related to iron uptake, such as the aerobactin and the *sitABCD* operons. We validated the association *in vivo* by deleting key components of the HPI in *E. coli* strains in two phylogenetic backgrounds, and found that virulence is correlated in *E. coli* with growth in the presence of various stressors including several antimicrobials, which hints at collateral sensitivities associated with intrinsic virulence. This study points to the major role of iron capture systems in the extra-intestinal virulence of the genus *Escherichia* and the collateral effects on cell growth of such systems.

## Introduction

Members of the *Escherichia* genus are both commensals of vertebrates^1^ and opportunistic pathogens^2^ involved in a wide range of intestinal and extra-intestinal infections. Apart from the *E. coli* species, it is composed of the cryptic *Escherichia* clades, and the *E. fergusonii* and *E. albertii* species. The latter taxons are rarely isolated in humans but are more frequently found in the environment and avian species where they can cause intestinal infections^3–5^. In humans, extra-intestinal infections represent a considerable burden^6^, with bloodstream infections (bacteraemia) being the most severe with a high attributable mortality of between 10-30%^7–10^. The regular increase over the last 20 years of *E. coli* bloodstream incidence^11^ and antibiotic resistance^12^ is particularly worrisome. The factors associated with high mortality are mainly linked to host conditions such as age, the presence of underlying diseases and to the portal of entry, with the urinary origin being more protective. These factors outweigh those directly attributable to the bacterial agent^7–9,13^.

Nevertheless, the use of animal models has shown a great variability in the intrinsic extra-intestinal virulence potential of natural *Escherichia* isolates. In a mouse model of sepsis where bacteria are inoculated subcutaneously, it has been clearly shown that the intrinsic virulence quantified by the number of animal deaths over the number of inoculated animals for a given strain is dependant on the number of virulence factors such as adhesins, toxins, protectins and iron capture systems^14–18^. One of the most relevant virulence factors is the so-called high-pathogenicity island (HPI), a 36 to 43 kb region encoding the siderophore yersiniabactin, a major bacterial iron uptake system19, which has also been shown to reduce the efficacy of innate immune cells to cause oxidative stress20. The deletion of the HPI results in a decrease in the intrinsic virulence in the mouse model in a strain-dependent manner^16,17,21^, indicating complex interactions between the genetic background of each strain and the HPI.

The limitation of these gene KO studies is that they target specific candidate genes. Recently, the development of new approaches in bacterial genome-wide association studies (GWAS)^22–25^ allows searching in an unbiased manner for genotypes associated with specific phenotypes such as drug resistance or virulence. In this context, we conducted a GWAS in 370 commensal and extra-intestinal pathogenic strains of *E. coli*, and related *Escherichia* clades, as well as *E. fergusonii* and *E. albertii*, representing the genus phylogenetic diversity, to search for traits associated with virulence in the mouse model of sepsis^26^. Most of the strains were isolated from a human host and are divided between commensals and extra-intestinal pathogenic. Most importantly, many (N=186) of these strains have been recently phenotyped across hundreds of growth conditions, including antibiotics and other chemical and physical stressors^27^. This data could then be used to find phenotype associations with virulence and to generate hypotheses on the function of genetic variants associated with the extra-intestinal virulence phenotype and potential collateral sensitivities associated with them.

## Results

### GWAS identifies the high-pathogenicity island as the strongest driver of the extra-intestinal virulence phenotype

We studied a 326 strain collection representative of the *E. coli* phylogenetic diversity, with strains belonging to phylogroups A (N=72), B1 (N=41), B2 (N=111), C (N=36), D (N=20), E (N=19), F (N=12) and G (N=15). To have a broader phylogenetic representation, we also included strains from *Escherichia* clades I to V (N=32) and the species *E. albertii* (N=7) and *E. fergusonii* (N=5)^28^. These strains encompass 170 commensal strains and 187 strains isolated in various extra-intestinal infections, mainly urinary tract infections and septicaemia^7,14,29–35^. To avoid any bias linked to host conditions, we assessed the strain virulence as its intrinsic extra-intestinal pathogenic potential using a well-calibrated mouse model of sepsis^14,26^, expressed as the number of killed mice over the 10 inoculated per strain. In accordance with previous data^14,26,36–38^, phylogroup B2 is the most associated with the virulence phenotype (2E^−9^ Wald test p-value, Figure 1A, Supplementary Table 1).

**Figure 1.**
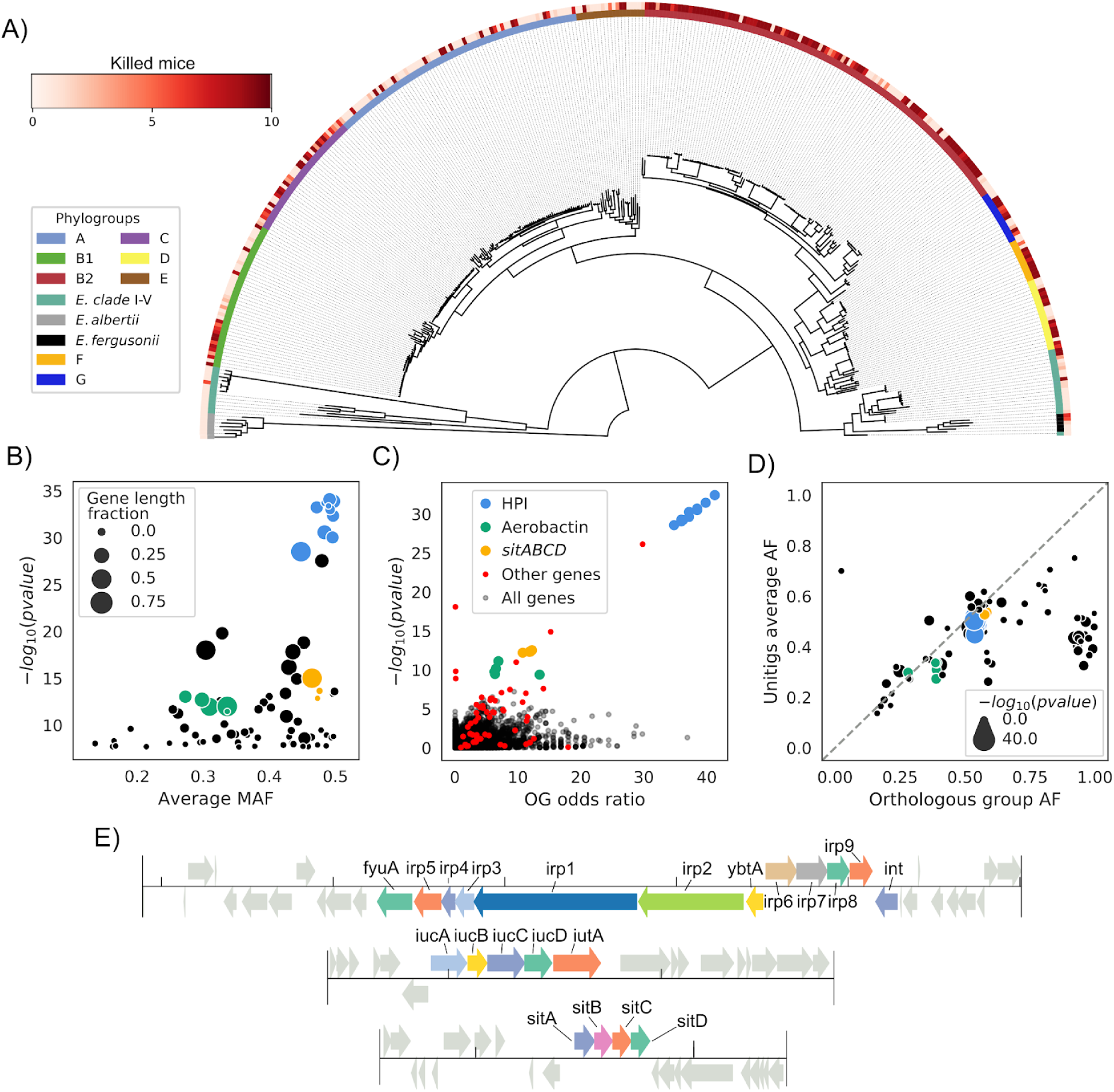
The HPI is strongly associated with the extra-intestinal virulence phenotype assessed in the mouse sepsis assay. A) Core genome phylogenetic tree of the *Escherichia* strains used in this study rooted on *E. albertii* strains. Outer ring reports virulence as the number of killed mice over the 10 inoculated per strain, inner ring the phylogroup or species each strain belongs to. B) Results of the unitigs association analysis: for each gene the minimum association p-value and average minimum allele frequency (MAF) across all mapped unitigs is reported. The gene length fraction is computed by dividing the total length of mapped unitigs by the length of the gene. C) Results of the gene presence/absence association analysis; each gene tested is represented. Genes from the unitigs association analysis are highlighted. OG: orthologous groups. D) Scatterplot of gene frequency versus frequency of associated unitigs; points on the diagonal indicate hits where the association is most likely due to a gene’s presence rather than a SNP. E) The structure of the HPI and of the aerobactin and *sitABCD* operons in strain IAI39; all associated genes are highlighted.

We used a bacterial GWAS method to associate unitigs to the virulence phenotype, allowing us to simultaneously test the contribution of core and accessory genome variation to pathogenicity^24^. It is generally understood that such methods require large sample sizes to have sufficient power, partly due to the need to break the long clonal frames typical of bacterial genomes; the appropriate sample size is also a function of the penetrance of the causal variants^23,39^. We ran simulations with an unrelated set of complete *E. coli* genomes and verified that our sample size was appropriate for variants with high penetrance (i.e. odds ratio above 5, Supplementary Figure 1, Methods). We reasoned that the genetic determinants of virulence are likely to have a relatively high penetrance, and that the strains used were genetically diverse, enough to break the clonal frame.

We uncovered a statistically significant association between 5,214 unitigs and the virulence phenotype, which were mapped back to 81 genes across the strains’ pangenome (Figure 1B, Supplementary Table 2, Methods). To understand whether the presence of these genes is directly associated with virulence or if its due to genetic variants such as SNPs we performed a separate association using genes’ presence absence patterns, showing that most genes have an odds ratio that far exceeds the required threshold we estimated from simulations, as well as low association p-value (Figure 1C). Furthermore, 48 out of 81 genes with at least an associated unitig mapped to them have a frequency across strains that is highly correlated with that of the associated unitigs (Figure 1D), indicating that it’s the presence of those genes to be associated with virulence. Genes belonging to the HPI had the lowest association p-value by far (<1E^−28^); the presence of two additional operons encoding for bacterial siderophores (aerobactin^40^ and *sitABCD41*) was also found to be associated with virulence (Figure 1E). We found that the HPI structure was highly conserved across the genomes that encode it (Supplementary Figure 2). We also observed that the distribution of known virulence factors didn’t match the virulence phenotype as closely as the HPI or the aerobactin and *sitABCD* operons, or had unitigs passing the association threshold (Supplementary Figure 3). The remaining 33 genes have a high frequency in the pangenome, indicating the presence of genetic variants such as SNPs present in core genes; the *mtfA* gene, which is involved in the regulation of carbohydrate metabolism^42^, had the lowest association p-value among those genes (1E^−14^).

### KO gene experiments validate the role of the HPI in the extra-intestinal phenotype

The studies on the role of the HPI in experimental virulence gave contrasting results according to the strains’ genetic background^17^. Among B2 phylogroup strains, HPI deletion in the 536 (ST127) strain did not have any effect in the mouse model of sepsis^43^ whereas this deletion in the NU14 (ST95) strain dramatically attenuated virulence^17^. Two strains from this study belonging to B2 phylogroup/ST141 (IAI51 and IAI52) deleted in *irp1* have attenuated virulence in the same model21. To have a broader view of the role of the HPI in various genetic backgrounds, we constructed *irp2* deletion gene mutants in two strains of phylogroup D (NILS46) and A (NILS9) belonging to STs (Sequence Types) frequently involved in human bacteraemia (ST69 and ST10, respectively)^44^. We first verified that the wild-type strains strongly produced yersiniabactin, whereas the *irp2* mutants did not (Figure 2A). We then tested them in the mouse sepsis model and saw an increase in survival for both mutated strains (log-rank test p-value < 0.0001 and 0.0217 for strain NILS9 and NILS46, respectively, Figure 2B, Supplementary Table 3) with no significant difference between the survival profiles for the two mutants (log-rank test p-value > 0.1).

**Figure 2.**
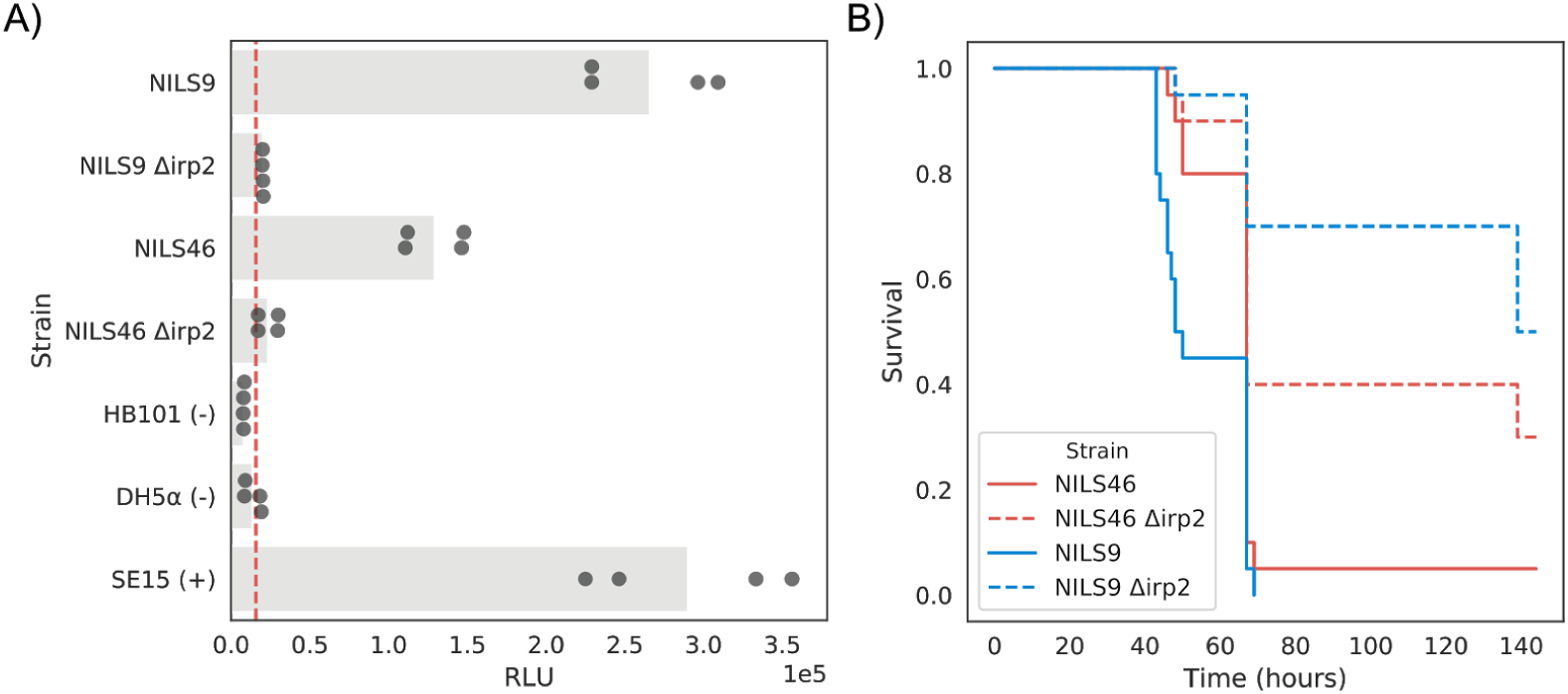
Phenotypic consequences of HPI deletion. A) Deletion of HPI leads to a decrease in production of yersiniabactin. Production of yersiniabactin is measured using a luciferase-based reporter (Methods). Strains marked with a “-” and “+” sign indicate a negative and positive control, respectively. The red dashed line indicates an arbitrary threshold for yersiniabactin production, derived from the average signal recorded from the negative controls plus two standard deviations. RLU, relative light units. B) Deletion of HPI leads to an increase in survival after infection. Survival curves for wild-type strains and the corresponding *irp2* deletion mutant, built after infection of 20 mice for each strain.

We have therefore validated *in vivo* the causal link between the HPI and the virulence phenotype detected by the means of an unbiased association approach, which demonstrates the power and accuracy of bacterial GWAS.

### High-throughput phenotypic data sheds light on HPI and other iron capture systems function

The main function encoded by the HPI cassette is iron scavenging through the expression of the siderophore yersiniabactin^21^, which has been previously validated in *E. coli* through knockout experiments^17^. The aerobactin operon also encodes an iron chelator^40^, while the *sitABCD* operon encodes a Mn^2+^/Fe^2+^ ion transporter^41^. In order to investigate other putative functions of these operons and their relationship with virulence, we leveraged a previously-generated high-throughput phenotypic screening in an *E. coli* strain panel that largely overlaps with the strains used here (186 over 370)^27^. We observed a relatively high correlation between growth profiles in certain conditions and both virulence and presence of the HPI, aerobactin and *sitABCD* operons (Figure 3A-D, Supplementary Table 4).

**Figure 3.**
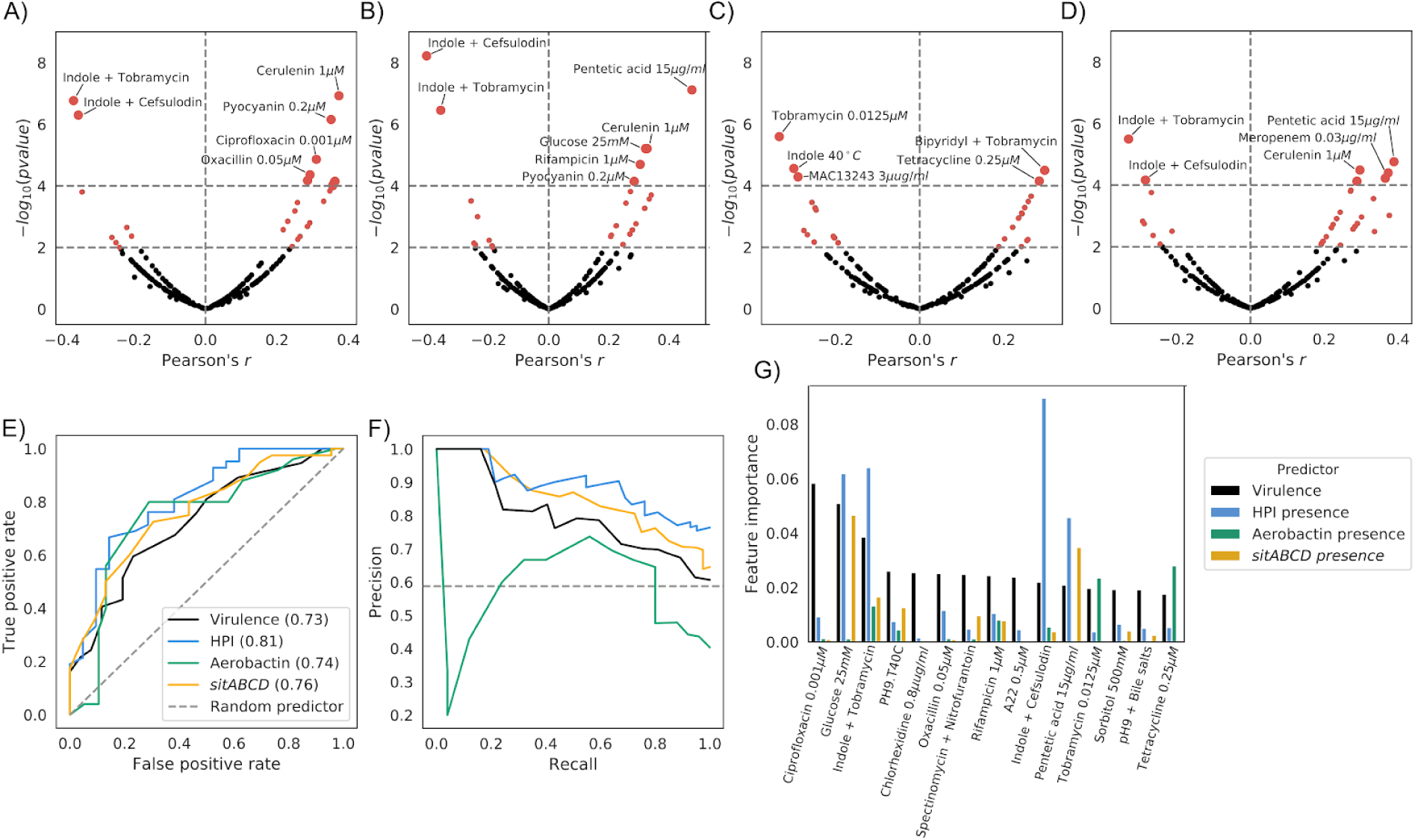
Growth profiles can predict virulence and presence of virulence factors. A-D) Volcano plots for the correlation between the strains’ growth profiles and: A) virulence levels, B) presence of the HPI, C) presence of aerobactin, and D) presence of *sitABCD*. E-F) Use of the strains’ growth profiles to build a predictor of virulence levels and presence of the three iron uptake systems. E) Receiver operating characteristic (ROC) curve and F) Precision-Recall curve for the four tested predictors. G) Feature importance for the predictors, showing the top 15 conditions contributing to the virulence levels predictor.

As expected, we found a positive correlation between growth on the iron-sequestering agent pentetic acid^45^ and both virulence and HPI/aerobactin/*sitABCD* presence (Pearson’s r: 0.36, 0.48, 0.23 and 0.39, respectively). We also found that growth in the presence of bipyridyl, an iron chelator, was positively correlated with the presence of aerobactin (bipyridyl + tobramycin, Pearson’s r: 0.30). We similarly observed a positive correlation between growth with pyocyanin, a redox-active phenazine compound being able to reduce Fe^3+^ to Fe^2+46^, and both HPI/aerobactin/*sitABCD* presence (Pearson’s r: 0.35, 0.28, 0.26 and 0.27 respectively). All growth conditions agree with the iron scavenging function of the three gene clusters and their importance for virulence.

Interestingly, we also found similarly strong positive correlations with growth on sub-inhibitory concentrations of several antibiotics, such as rifampicin, ciprofloxacin, tetracycline and ß-lactams such as amoxicillin, oxacillin and meropenem, as well as other antimicrobial agents such as cerulenin and colicin. On the other hand we found that growth in presence of indole at 2 mM either in association with sub-inhibitory concentrations of cefsulodin and tobramycin, or alone at 40°C was negatively correlated with both virulence and HPI/aerobactin/*sitABCD* presence. Similar negative correlation was observed with aerobactin presence and the MAC13243 compound that increases outer membrane permeability^47^. These correlations might be due to the presence/absence of acquired resistance alleles and/or genes that are strongly associated with pathogenic strains, or might point to the role of iron homeostasis in intrinsic resistance to antibiotics^48^. To investigate these two hypotheses, we focused on tetracycline resistance, a common occurrence in the genus^33,37,49^, and for which resistance genes can be easily found through sequence homology (Methods). We measured the correlation between the presence of tetracycline resistant genes, found in 26.8% of the strains, and virulence (Pearson’s r: 0.16), as well with the presence of either of the three iron capture systems (Pearson’s r: 0.21, 0.33 and 0.24 for HPI, aerobactin and *sitABCD*, respectively), which we found to be all comparable with the direct correlation between growth on sub-inhibitory concentration of tetracycline and the presence of resistance genes (Pearson’s r: 0.4). As such, both scenarios seem plausible to explain the relationship between growth in presence of tetracycline and both virulence and presence of HPI/aerobactin/*sitABCD*.

Given the relatively large number of conditions correlated with both virulence and presence of iron uptake systems, we tested whether these features could be predicted from growth profiles. We used the commonly-used random forests machine learning algorithm with appropriate partitioning of input data to tune hyperparameters and reduce overfitting, leading to four classifiers for virulence and presence of the HPI, aerobactin and *sitABCD* operons with high predictive power, with the exception of aerobactin (Figure 3E and 3F and Methods). We noted that prediction of the gene clusters presence performs slightly better than virulence, possibly reflecting the complex nature of the latter phenotype. As expected, we found that conditions with relatively high correlation with each feature have a higher weight across classifiers (Figure 3G, Supplementary Table 5), which suggests that a subset of phenotypic tests might be sufficient to classify pathogenic strains. These results show how phenotypic data can be used to generate hypotheses for the function of virulence factors.

## Discussion

With the steady decline in the price of genomic sequencing and the increasing availability of molecular and phenotypic data for bacterial isolates, it has finally become possible to use statistical genomics approaches such as GWAS to uncover the genetic determinants of relevant phenotypes. Such approaches have the advantage of being unbiased, and can then be used to confirm previous targeted findings and potentially uncover new factors, given sufficient statistical power. The accumulation of other molecular and phenotypic data can on the other hand uncover variables correlated with phenotype, which can be used to generate testable hypotheses on the function of genomic hits and potential collateral sensitivities associated with them. Given the rise of both *E. coli* extra-intestinal infections and antimicrobial resistance, we reasoned that the intrinsic virulence assessed in a calibrated mouse model of sepsis^14,26^ is a phenotype worth exploring with such an unbiased approach.

We were able to confirm earlier reports on the importance of the HPI in extra-intestinal virulence^17–19,21,50,51^, which showed the strongest signal in both the unitigs and accessory genome association analysis, and whose importance was validated *in vitro* and in an *in vivo*. The distribution of the HPI within the species resulting from multiple horizontal gene transfers via homologous recombination^52^ has probably facilitated its identification using GWAS. We associated additional genetic factors to intrinsic virulence, such as the aerobactin and *sitABCD* operons, both related to iron scavenging together with the HPI. We also found mutations in the *mtfA* core gene to be associated with virulence, although its role in virulence remains elusive. Additional factors might have been overlooked by this analysis, due to the relatively small sample size; we however estimate that those putative additional factors might have a relatively low penetrance, based on our simulations in an independent dataset. As sequencing of bacterial isolates is becoming more common in clinical settings^53–55^, we expect to be able to uncover these additional genetic factors in future studies.

The association between both the intrinsic virulence phenotype and the presence of the virulence factors - such as the HPI - and previously collected growth data allowed us to generate testable hypotheses on mechanism of pathogenesis and putative additional functions of these factors. In particular we observed a relatively strong correlation between growth on various antimicrobial agents and both virulence and HPI/aerobactin/*sitABCD* presence, which seems to confirm the pressure to acquire resistance for these isolates, as exemplified for tetracycline resistance genes, but also raises the question about the potential role of iron homeostasis on antimicrobial resistance^48^. As we observed several classes of antibiotics with different molecular targets, it is possible that this is due to a pleiotropic effect. *E. coli* Fur mutants, a transcriptional regulator that represses iron uptake systems, which accumulate high level of intracellular iron, have been shown to increase resistance to quinolones, aminoglycoside, tetracycline, rifampicin and amoxicillin^56^. Cell envelope permeability can also be modified in response to the presence of iron via two-component systems, rendering the cell more resistant^48^. Sub-lethal doses of antibiotics have also been shown to behave as a stressor for the cells that engage a general core hormetic stress response consisting of increased energy production and translation, allowing growth when nutrients are available despite the constant presence of the stressor^57^. When the resources are instead limited, cell growth is stopped. It can be hypothesised that iron is part of these limiting nutrients and that iron capture systems allow coping with the stressors, i.e. the antibiotics.

The negative correlation between virulence and iron capture systems and growth profiles in the presence of indole associated with stress conditions such as sub-lethal doses of antibiotics or high temperature but not indole alone, points however to the possible deleterious role of iron in such conditions. In lysogeny broth, sub-lethal concentrations of antibiotics increase the endogenous production of indole^57^ and, at very high concentration (*e.g.* 5 mM), indole induces the production of reactive oxygen species and is toxic for the cells^58^. A vicious circle is rapidly established as antibiotics increase the production of indole^57^, which in turn destabilises the membrane^58^, further increasing the penetration of the antibiotics. This toxicity has been shown to be partly iron mediated due to the Fenton reaction, the deletion of TonB, an iron transporter, increasing resistance to the antibiotic^59^. Interestingly, tobramycin is the antibiotic involved in this negative correlation (Fig. 3A to D). Sub-lethal doses of this aminoglycoside lead to an increase of reactive oxygen species in the bacterial cell in relation to intra-cellular iron and Fenton reaction^60^. Thus, cells with increased import of extracellular iron might be more sensitive to sub-lethal doses of antibiotics, suggesting the potential for collateral sensitivities related to both intrinsic virulence and the presence of the iron uptake systems. Altogether, these data bring new light on the “liaisons dangereuses” between iron and antibiotics that could potentially be targeted^48^. More generally, they show that the presence of iron capturing systems can be either advantageous or disadvantageous, depending on the growth conditions.

In conclusion, we showed the power of bacterial GWAS to identify major virulence determinants in bacteria. Within the *Escherichia* genus, iron capture systems are the main predictors of the intrinsic extra-intestinal virulence. Furthermore, this analysis demonstrates how a data-centric approach can increase our knowledge of complex bacterial phenotypes and guide further empirical work on gene function and its relationship to intrinsic virulence.

## Materials and methods

### Strains used

The full list of the 370 strains used in the association analysis, together with their main characteristics is reported in Supplementary Table 1. These strains belong to various published collections: ECOR (N=71)^29^, IAI (n=81)^14^, NILS (N=82)^31^, Septicoli (N=39)^10^, ROAR (N=30)^32^, Guyana (N=12)^30^, Coliville (N=8)^33^, FN (N=6)^34^, COLIRED (N=3)^35^, COLIBAFI (N=2)^7^, correspond to archetypal strains (N=7) or are miscellaneous strains from our personal collections (N=29). The isolation host is predominantly humans (N=291), followed by animals (N=72) and isolated from the environment (N=6). One hundred and seventy strains were commensal whereas five and 187 were responsible of intestinal and extra-intestinal infections, respectively. The genomes of 295 strains were previously available, while the remaining 75 were sequenced as part of this work by Illumina technology as described previously^35^. The genome sequences of all strains are available through Figshare^61^.

The construction of the *irp2* deletion mutants of the NILS9 and NILS46 strains was achieved following a strategy adapted from Datsenko and Wanner^62^. Primers used in the study are listed in Supplementary Table 6. In brief, primers used for gene disruption included 44-46 nucleotide homology extensions to the 5’- and 3’ regions of the target gene, respectively, and additional 20 nucleotides of priming sequence for amplification of the resistance cassette on the template plasmids pKD4. The PCR product was then transformed into strains carrying the helper plasmid pKOBEG expressing the lambda red recombinase under control of an arabinose-inducible promoter^63^. Kanamycin resistant transformants were selected and further screened for correct integration of the resistance marker by PCR. For elimination of the antibiotic resistance gene, helper plasmid pCP20 was used according to the published protocol. PCR followed by Sanger sequencing of the mutants were performed to verify the deletion and the presence of the expected scar.

### Yersiniabactin detection assay

Production of the siderophore yersiniabactin was detected and quantified using a luciferase reporter assay as described elsewhere^17,64^. Briefly, bacterial strains were cultivated in NBD medium for 24 hours at 37°C. Next, bacteria were pelleted by centrifugation and the supernatant was added to the indicator strain WR 1542 harbouring plasmid pACYC5.3L. All the genes necessary for yersiniabactin uptake are located on the plasmid pACYC5.3L, i.e. *irp6, irp7, irp8, fyuA, ybtA*. Furthermore, this plasmid is equipped with a fusion of the *fyuA* promoter region with the luciferase reporter gene. The amount of yersiniabactin can be quantified semi-quantitatively, as yersiniabactin-dependant upregulation of *fyuA* expression is determined by luciferase activity of the *fyuA-luc* reporter fusion.

### Mouse virulence assay

Ten female mice OF1 of 14-16 g (4 week-old) from Charles River® (L’Arbresle, France) received a subcutaneous injection of 0.2 ml of bacterial suspension in the neck (2·10^8^ colony forming unit). Time to death was recorded during the following 7 days. Mice surviving more than 7 days were considered cured and sacrificed1^14^, In each experiment, the *E. coli* CFT073 strain was used as a positive control killing all the inoculated mice whereas the *E. coli* K-12 MG1655 strain was used as a negative control for which all the inoculated mice survive^26^. For the mutant assays, 20 mice per strain were used to obtain statistical relevant data. The data was analysed using the lifeline package v0.21.0^65^.

### Association analysis

All genome-wide association analysis were carried out using pyseer, version v1.3.4^24^. All input genomes were re-annotated using prokka, version v1.14.566, to ensure uniform gene calls and excluding contigs whose size was above 200 base pairs. The core genome phylogenetic tree was generated using ParSNP67 to generate the core genome alignment and gubbins v2.3.5^68^ to generate the phylogenetic tree. The strain’s pangenome was estimated using roary v3.13.0^69^. Unitigs distributions from the input genome assemblies were computed using unitig-counter v1.0.5. The association between both unitigs and pangenome and phenotype (expressed as number of mice killed post-infection) was carried out using the FastLMM^70^ linear mixed-model implemented in pyseer, using a kinship matrix derived from the phylogenetic tree as population structure. For both association analysis we used the number of unique presence/absence patterns to derive an appropriate p-value threshold for the association likelihood ratio test (2.16E^−08^ and 5.45E^−06^ for the unitigs and pangenome analysis, respectively). Unitigs significantly associated with the phenotype were mapped back to each input genome using bwa mem v0.7.17-r1188^71^ and betools v.2.29.2^72^, using the pangenome analysis to collapse gene hits to individual groups of orthologs. A sample protein sequence for each groups of orthologs where at least one unitig with size 20 or higher was mapped was extracted giving priority to strain IAI39 when available, given it was the only strain with a complete genome available^73^; those sample sequences where used to search for homologs in the uniref50 database from uniprot^74^ using blast v2.9.0^75^. Each group of orthologs was then given a gene name using both available literature information and the results of the homology search.

### Power simulations

Statistical power was estimated using an unrelated set of 548 complete *E. coli* genomes downloaded from NCBI RefSeq using ncbi-genome-download v0.2.9 on May 24th 2018. Each genome was subject to the same processing as the actual ones used in the real analysis (re-annotation, phylogenetic tree construction, pangenome estimation). The gene presence/absence patterns were used to run the simulations, in a similar way as described in the original SEER implementation^23^. Briefly, for each sample size, a random subset of strains was selected, and the likelihood ratio test p-value threshold was estimated by counting the number of unique gene presence/absence patterns in the reduced roary matrix. For each odds ratio tested, a binary case-control phenotype vector was constructed for the strains subset using the following formulae:

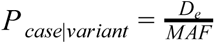

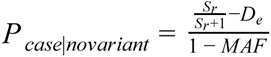

Were *S*_*r*_ is the ratio of case/controls (set at 1 in these simulations), *MAF* as the minimum allele frequency of the target gene in the strains subset, and *D*_*e*_ the number of cases. pyseer’s LMM model was then applied to the presence/absence vector of the target gene and the likelihood ratio test p-value was compared with the empirical threshold. The randomization was repeated 100 times and power was defined as the proportion of randomizations for each sample size and odds ratio whose p-value was below the threshold. The *pks2* and *fabG* genes were used as gene targets in the simulations, and both gave very similar results.

### Correlations with growth profiles

The previously generated phenotypic data^27^ for 186 over 370 strains were used to compute correlations with both the number of mice killed after infection and presence/absence of the associated virulence factors. The data was downloaded from the ecoref website (https://evocellnet.github.io/ecoref/download/) and the pearson correlation with the s-scores was computed together with the correlation p-value. Prediction of tetracycline resistance was carried out using staramr v0.7.1 with the ResFinder database^76^. Four predictors, one for virulence (number of killed mice post-infection) and one for presence of the HPI, aerobactin and the *sitABCD* operon were built using the random forest classifier algorithm implemented in scikit-learn v.022.0^77^, using the s-scores as predictors. The input was column imputed, and 33% of the observations were kept as a test dataset, using a “stratified shuffle split” strategy. The remainder was used to train the classifier, using a grid search to select the number of trees and the maximum number of features used, through 10 rounds of stratified shuffle split with validation set size of 33% the training set and using the F1 measure as score. The performance of the classifiers on the test set were assessed by computing the area under the receiver operating characteristic curve (ROC-curve).

## Code and data availability

All input data and code used to run the analysis and generate the plots is available online at https://github.com/mgalardini/2018_ecoli_pathogenicity. Code is mostly based on the Python programming language and the following libraries: numpy v1.17.3^78^, scipy v1.4.0^79^, biopython v1.75^80,81^, pandas v0.25.3^82^, pybedtools v0.8.0^83^, dendropy 4.4.0^84^, ete3 v3.1.1^85^, statsmodels v0.10.2^86^, matplotlib v3.1.2^87^, seaborn v0.9.0^88^, jupyterlab v1.2.4^89^ and snakemake v5.8.2^90^.

## Ethics statement

All animal experimentations were conducted following European (Directive 2010/63/EU on the protection of animals used for scientific purposes) and national recommendations (French Ministry of Agriculture and French Veterinary Services, accreditation A 75-18-05). The protocol was approved by the Animal Welfare Committee of the Veterinary Faculty in Lugo, University of Santiago de Compostela (AE-LU-002/12/INV MED.02/OUTROS 04).

## Acknowledgements

We are grateful to Ivan Matic for discussion on the effect of indole. This work was partially supported by the “Fondation pour la Recherche Médicale” (Equipe FRM 2016, grant number DEQ20161136698).

## Supplementary Figures

**Supplementary Figure 1.**
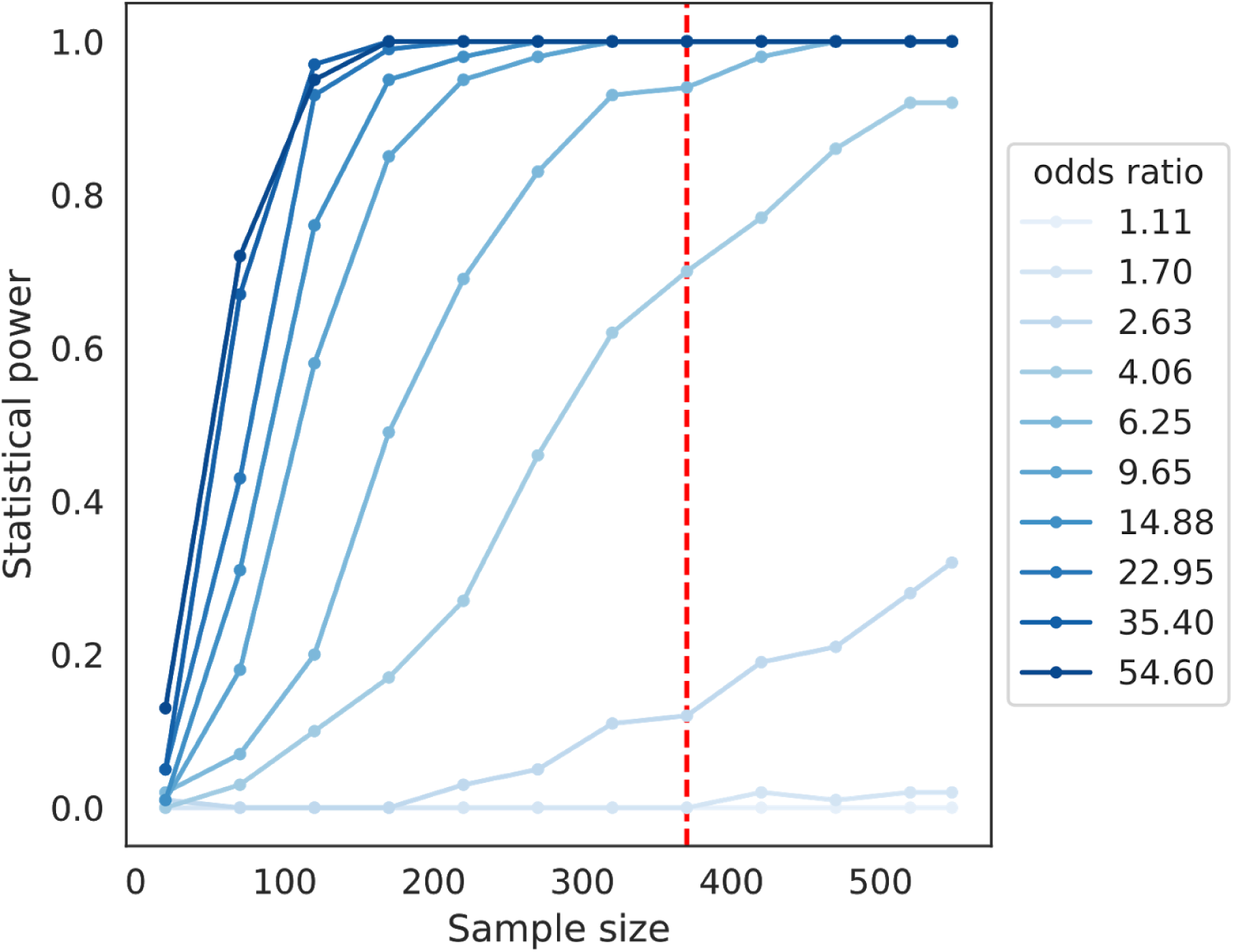
Simulations of statistical power on an unrelated set of complete *E. coli* genomes, using the *pks2* gene as target. The dotted red line indicates the sample size used in the actual analysis.

**Supplementary Figure 2.**
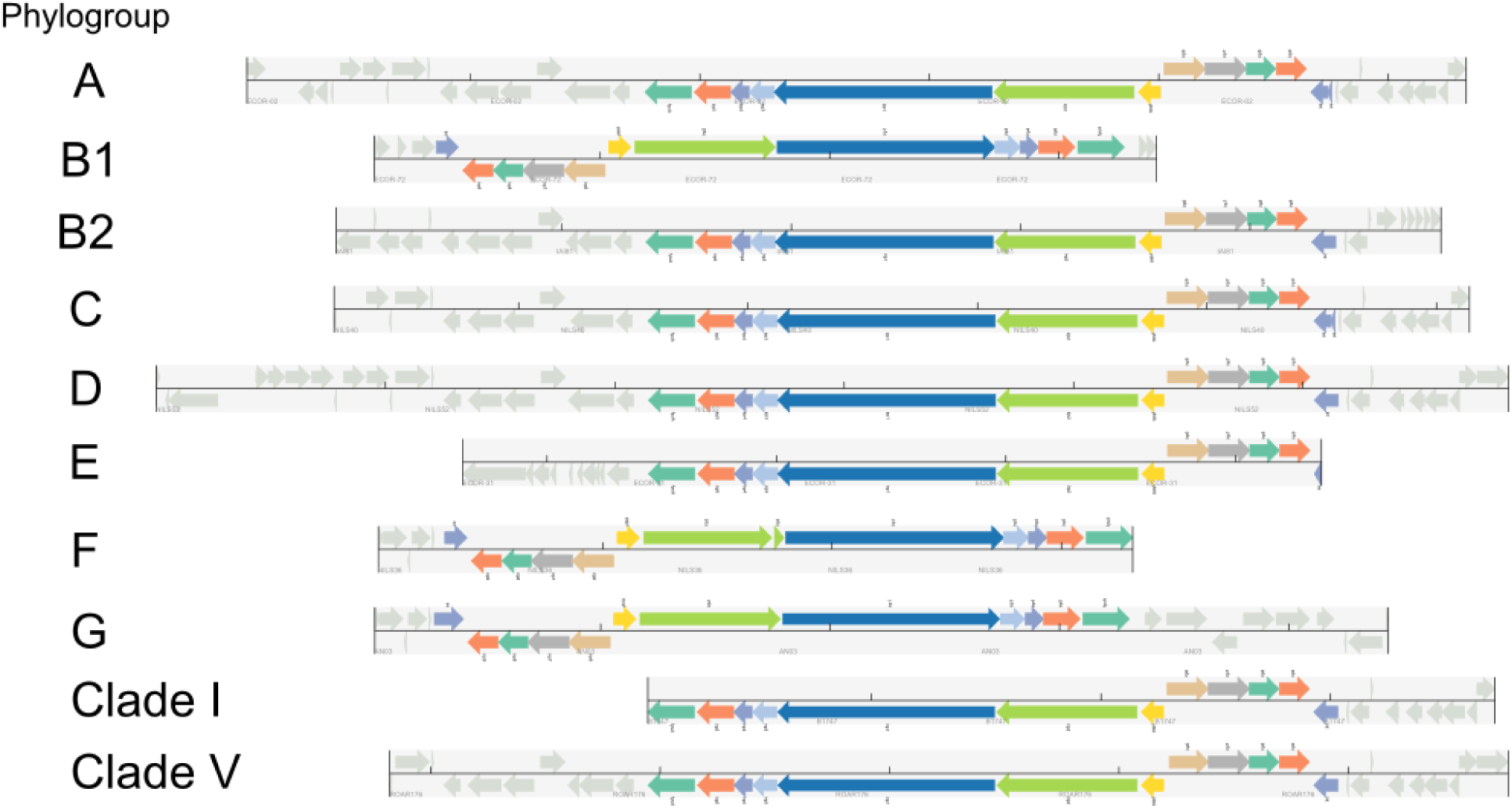
HPI structure conservation across strains. One strain per phylogroup or species is shown, using the same color scheme as Figure 1E for each gene.

**Supplementary Figure 3.**
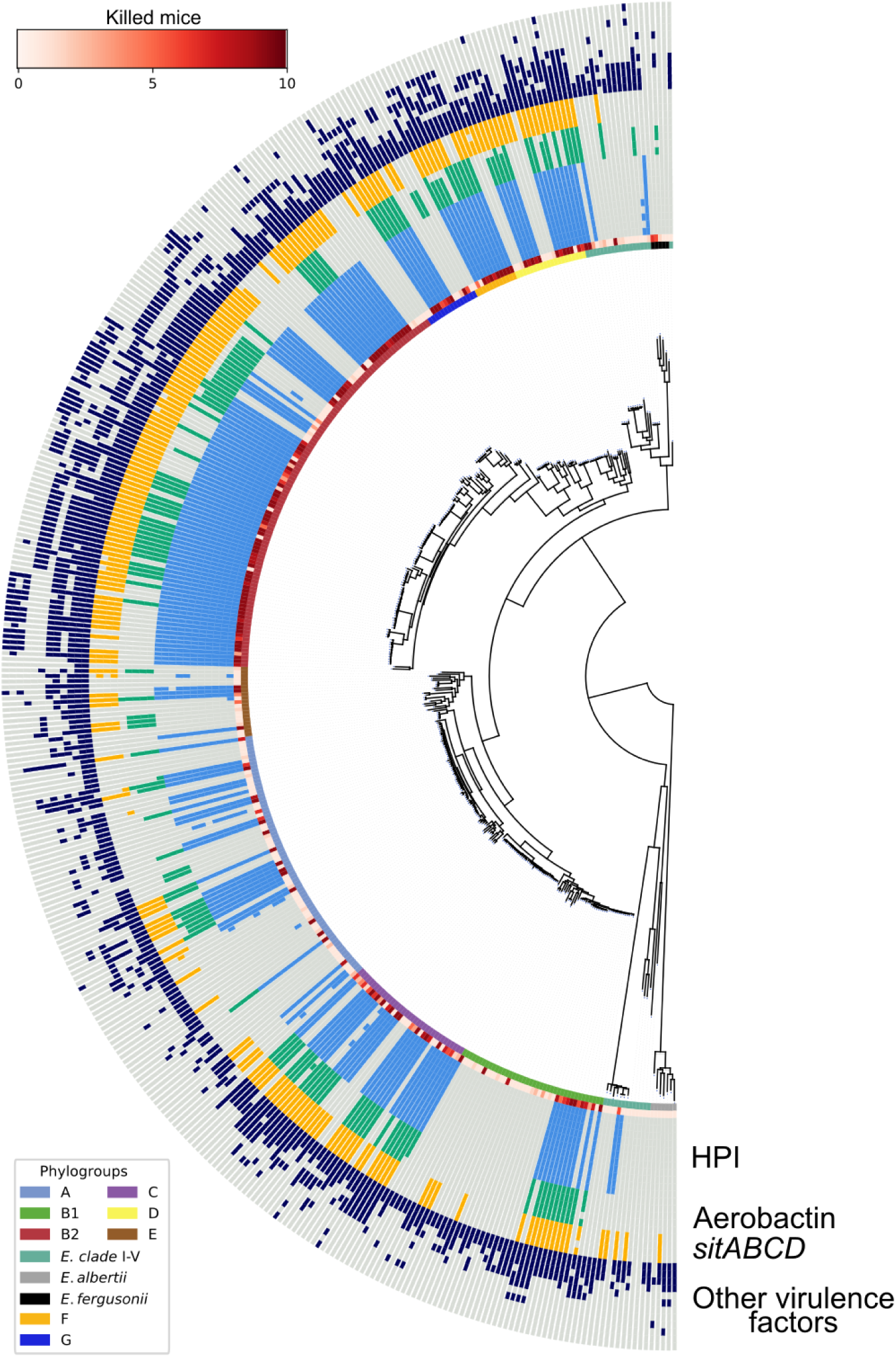
Presence/absence patterns of known virulence factors. Solid color indicates presence, light grey indicates absence. Phenotypes (number of killed mice) and phylogroup or species of each strain are reported as in Figure 1A. “Other virulence factors” are (from inside the ring towards the outside): *sfaD*, *sfaE*, *ompT*, *traT*, *hra2*, *papC*, *iha*, *ireA*, *neuC*, *hlyC*, *clbQ* and *cnf1*.

## Supplementary Information

**Supplementary Table 1:** Strains’ information, including virulence phenotype

**Supplementary Table 2:** Summary of the 81 genes with at least one mapped unitig

**Supplementary Table 3:** Survival analysis for NILS9 and NILS46 wild-type and HPI mutants

**Supplementary Table 4:** Correlation between growth on stress conditions (s-scores) and both virulence and presence of the HPI

**Supplementary Table 5:** Feature importance for each growth condition in the random forests predictor for virulence and HPI presence

**Supplementary Table 6:** List of PCR primers used in this study

